# The Role of C-to-U RNA Editing in Human Biodiversity

**DOI:** 10.1101/2023.07.31.550344

**Authors:** Melissa Van Norden, Zackary Falls, Sapan Mandloi, Brahm Segal, Bora Baysal, Ram Samudrala, Peter L. Elkin

## Abstract

Intra-organism biodiversity is thought to arise from epigenetic modification of our constituent genes and post-translational modifications after mRNA is translated into proteins. We have found that post-transcriptional modification, also known as RNA editing, is also responsible for a significant amount of our biodiversity, substantively expanding this story. The APOBEC (apolipoprotein B mRNA editing catalytic polypeptide-like) family RNA editing enzymes APOBEC3A and APOBEC3G catalyze the deamination of cytosines to uracils (C>U) in specific stem-loop structures.^1,2^ We used RNAsee (RNA site editing evaluation), a tool developed to predict the locations of APOBEC3A/G RNA editing sites, to determine whether known single nucleotide polymorphisms (SNPs) in DNA could be replicated in RNA via RNA editing. About 4.5% of non-synonymous SNPs which result in C>U changes in RNA, and about 5.4% of such SNPs labelled as pathogenic, were identified as probable sites for APOBEC3A/G editing. This suggests that the variant proteins created by these DNA mutations may also be created by transient RNA editing, with the potential to affect human health. Those SNPs identified as potential APOBEC3A/G-mediated RNA editing sites were disproportionately associated with cardiovascular diseases, digestive system diseases, and musculoskeletal diseases. Future work should focus on common sites of RNA editing, any variant proteins created by these RNA editing sites, and the effects of these variants on protein diversity and human health. Classically, our biodiversity is thought to come from our constitutive genetics, epigenetic phenomenon, transcriptional differences, and post-translational modification of proteins. Here, we have shown evidence that RNA editing, often stimulated by environmental factors, could account for a significant degree of the protein biodiversity leading to human disease. In an era where worries about our changing environment are ever increasing, from the warming of our climate to the emergence of new diseases to the infiltration of microplastics and pollutants into our bodies, understanding how environmentally sensitive mechanisms like RNA editing affect our own cells is essential.

## Introduction

Our biodiversity has long been thought to come from alternative splicing and post-translational modification of the thousands of proteins encoded in the human genome. There are about 20,300 protein-encoding genes in the genome, of which 19,267 have an approved HUGO gene name (as of 04/07/23).^3,4^ However, about 70,000 proteins result from splice variants, and thousands more could result from post-translational modifications.^4^ The expansion of proteoforms from genes is multifactorial, we hypothesize, and it could be due to mechanisms beyond post-translational modifications. In this paper, we explore the ability of post-transcriptional modifications, or RNA editing, to contribute to human biodiversity.

RNA editing has been described as “any site-specific alteration in an RNA sequence that could have been copied from the template, excluding changes due to processes such as RNA splicing and polyadenylation.”^5^ There are two well-studied families of RNA editing enzymes which catalyze single nucleotide substitutions.^6^ The ADAR (adenosine deaminase acting on RNA) family deaminates adenosine to guanosine-analog inosine (A>I) in specific double-stranded RNA contexts.^7^ Similarly, specific APOBEC (apolipoprotein B mRNA editing catalytic polypeptide-like) family enzymes deaminate cytosine to uracil (C>U) in single-stranded contexts.^1,2^

Two such APOBEC-family enzymes are APOBEC3A and APOBEC3G. The ability of APOBEC3 enzymes to edit RNA is a relatively recent discovery, starting in 2015 with a paper by Sharma et al.^8^ For a long time, these enzymes were primarily known for their ability to edit single-stranded DNA (ssDNA) produced by viruses such as HIV (APOBEC3G) and parvovirus (APOBEC3A).^1^ Perhaps because of this, known APOBEC3-mediated RNA editing sites are rare. In a dataset taken from a paper by Asaoka et al., only about 0.05% of bases across 2343 genes were considered APOBEC3A/G RNA editing sites.^9^ We contend that, beyond their effects on viruses, these enzymes may also affect human health via the editing of healthy human mRNA and subsequent creation of protein variants.

Previous work has shown that APOBEC3A/G cytosine deaminases preferentially target RNA and ssDNA substrates with stem-loop structures.^2,10^ Specifically, an optimal target contains a tri- or tetraloop, with the edited cytosine at the 3’ end of the loop and a pyrimidine 5’ to it (Figure 1A).^2,10^ Specific cytosines for which APOBEC3 enzymes have high affinity, such as c.136 in SDHB, have previously been observed to undergo RNA editing even in normal physiological circumstances.^8,11^ Additionally, APOBEC3-mediated RNA editing activity is known to transiently increase in monocytes, peripheral blood cells, and blood-brain barrier cells upon environmental interferon exposure.^8,12-15^ Thousands of cytosines are known targets of APOBEC3-mediated editing, editing at many of which result in non-synonymous or nonsense mutations.^8,9,12^ Therefore, variant proteins that impact human health could be created by APOBEC3-mediated RNA editing at high-affinity sites and at an increased rate during and following tissue inflammation.

**Figure 1.**
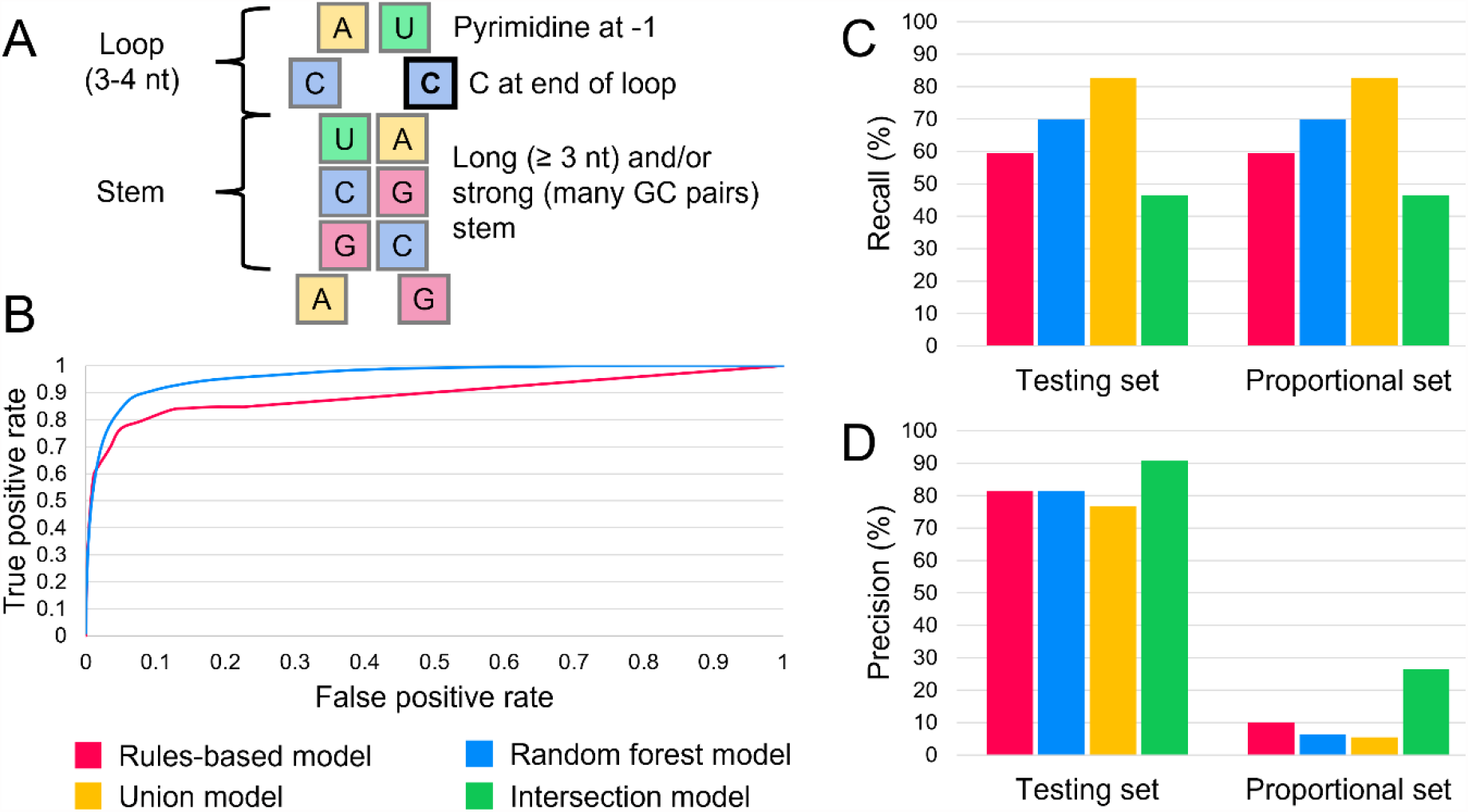
Benchmarking results of RNAsee. (A) APOBEC3A and APOBEC3G preferentially edit cytosines in stem-loop structures. The edited cytosine is outlined in black. (B) When tested on the proportional set, the random forest model had an AUROC of 0.962, and the rules-based model had an AUROC of 0.892. The ROC curves overlapped at higher thresholds (low false positive rate), but the random forest model greatly outperformed the rules-based model at lower thresholds. The recall (C) and precision (D) of the two primary models and two consensus models were assessed on the testing set (editing:non-editing site ratio of 1:3) and the proportional set (editing:non-editing site ratio of 1:468). Because both sets contained the same positive sites, recall was the same on both sets. The intersection model was the most precise on both sets, making it useful for selective studies; the union model had the greatest recall, so it can be used to survey potential editing sites more broadly.

There is a growing body of evidence supporting the influence of C>U RNA editing on human health. APOBEC3-mediated RNA editing sites have been found in genes with known relationships to neoplasms, hypertension, and nervous system disorders, including amyotrphic lateral sclerosis (ALS), Alzheimer’s, Huntington’s, and Parkinson’s disease.^8,12^ Correlation between APOBEC3-mediated C>U RNA editing and diseases like epilepsy and sporadic Creutzfeldt-Jakobs disease (sCJD) has been found in mice, and a link between RNA editing and autoantigen creation in autoimmune diseases like systemic lupus erythematosus (SLE) has been suggested in humans.^16-18^

However, this area of research is still relatively young. Most articles and tools on RNA editing still focus on ADAR-mediated A>I editing. For instance, the publicly available RADAR database catalogues A>I RNA editing sites with manual annotations, and this tool has been used alongside data from The Cancer Genome Atlas to identify protein variants that may affect tumor cell viability and drug sensitivity.^19,20^ Although at least one study attempted to collate known APOBEC3A/G editing sites, no comprehensive database like RADAR yet exists for C>U editing events, and studies on APOBEC3 editing tend to remain narrowly focused on one or two diseases at a time.^9^

To address this gap in knowledge, we have developed RNAsee (RNA site editing evaluation), a program that combines machine learning and rules-based methods to predict APOBEC3A/G mediated RNA editing sites in transcripts of human genes.^21^ In this paper, we will compare this list of predicted APOBEC3A/G-mediated C>U editing sites with publicly available data on human protein variants from the CinVar database, particularly pathogenic variants, to help open the conversation on the extent to which C>U RNA editing may contribute to human disease.

## Results

### Performance of RNAsee

To assess the performance of RNAsee in predicting APOBEC3A/G editing sites, we benchmarked the random forest and rules-based models on the proportional set, which contains a 468:1 ratio of non-editing to editing sites (see Performance benchmark for RNAsee). First, we calculated area under the receiver operator characteristic (AUROC) metrics for the rules-based and random forest models (Figure 1B). The AUROC of the random forest model was 0.962, which is higher than the rules-based model’s AUROC of 0.892. However, the two curves overlapped at the highest thresholds (lowest false positive rates), including the points at which the thresholds for the rules-based and random forest models were set.

Using a score threshold of >= 10 for the rules-based model and probability threshold of > 0.5 for the random forest model, we found recall and precision metrics for the rules-based and random forest models, plus two consensus models created using set operations (intersection and union) on the output of these two models. These metrics were calculated for the testing and proportional sets, which differed in the number of non-editing sites they included (see Performance benchmark for RNAsee). The recall of each model is shown in Figure 1C, and precision is shown in Figure 1D.

### Identification of possible editing sites

To start, we needed a set of nucleotide polymorphisms. We extracted an annotated set of DNA mutations from the ClinVar database.^22^ We pared this set down to only include single nucleotide polymorphisms (SNPs) which were exonic and non-synonymous. Then, we examined the relative frequency of different polymorphisms (Figure 2A and 2B).

**Figure 2.**
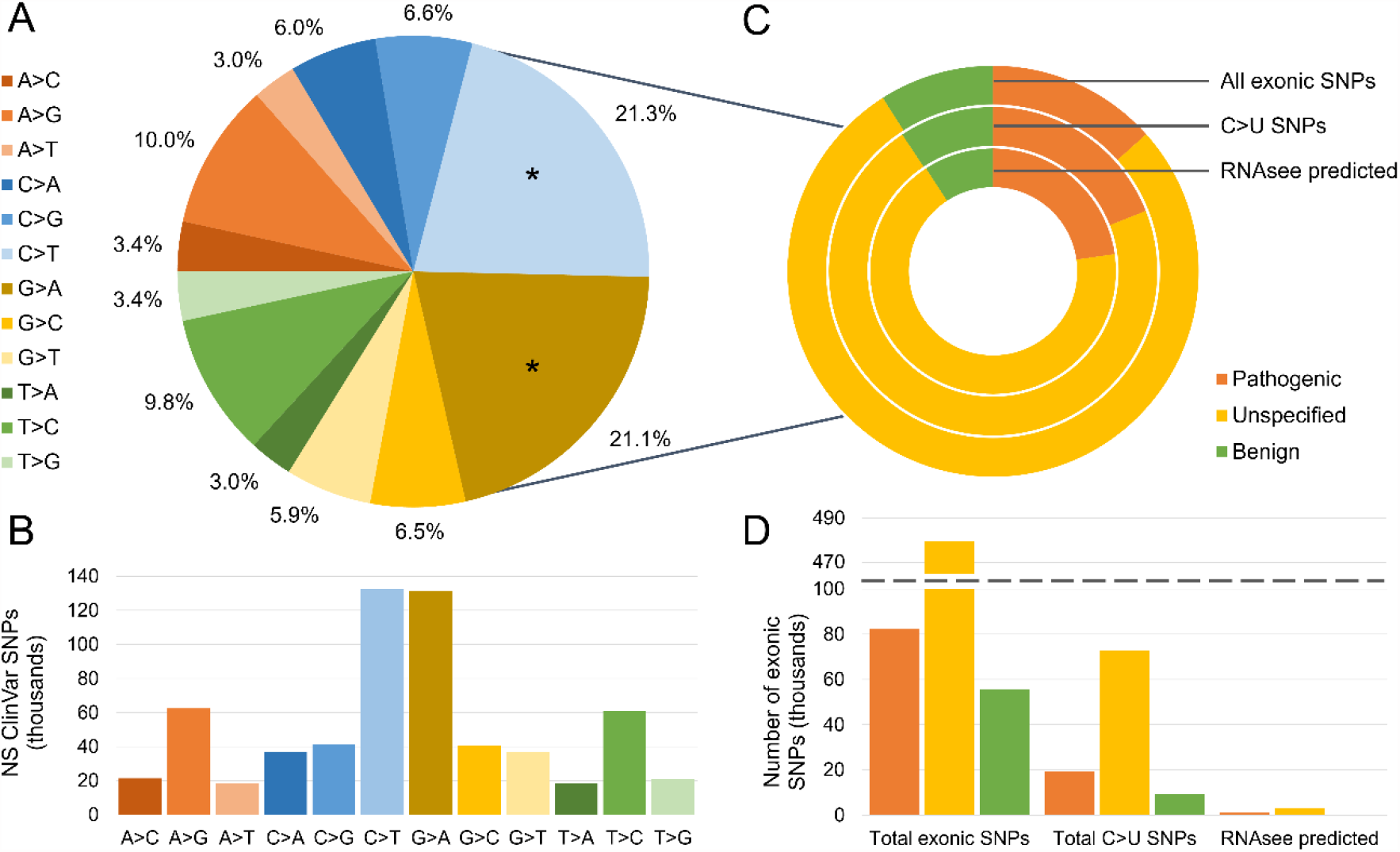
Over two-fifths of non-synonymous SNPs in the ClinVar database may result from C>U DNA editing. (A) The recorded nucleotide changes of all non-synonymous (NS) SNPs in the ClinVar database were counted. Like-to-like or non-specific changes were excluded. C>T (21.3%) and G>A (21.1%) mutations may be associated with C>U mutations when transcribed into RNA, depending on which strand is the template; the slices representing these changes are asterisked. The raw number of each type of NS SNP is plotted in (B). (C) Each ClinVar SNP is also associated with a pathogenicity label. These were sorted into three bins: “pathogenic,” “unspecified,” and “benign.” The proportions of each label among all exonic SNPs, all SNPs associated with C>U RNA changes, and all SNPs returned as possible APOBEC3 editing sites by RNASee are shown. The raw number of SNPs per site are shown in (D). 22.7% of the potential editing sites returned by RNAsee were labelled as likely pathogenic or pathogenic, whereas only 9.2% were labelled as likely benign or benign. This suggests that C>U RNA editing has a substantial possibility of negatively influencing human health.

Both C>T and G>A SNPs may result in C>U changes in RNA, depending on which DNA strand is transcribed. Among 622,622 non-synonymous SNPs, C>T (21.3%) and G>A (21.1%) were by far the most frequently observed polymorphisms. Both of these types of SNPs may result in C>U changes in RNA, depending on which DNA strand is transcribed. Therefore, to determine which of these SNPs could correspond with C>U editing, we needed to refer to the RNA sequences.

We examined coding sequence files for each gene with an SNP. Entries in genes with no good coding sequence file found were excluded, leaving 617,363 SNPs across 9228 genes. This set is the set of “all exonic SNPs”. Of these, 101,565 (16.5%) were associated with a C>U RNA change and were included in the set of “all C>U SNPs.” We used the most sensitive RNAsee model, the union model, to analyze each cytosine in that set. RNAsee returned 4600 SNPs, 4.5% of all C>U SNPs, as potential APOBEC3A/G-mediated RNA editing sites. These SNPs were included in the “RNAsee predicted” set. Of these 4600 sites, 62 (1.34%) are known APOBEC3-mediated RNA editing sites included in the Asaoka et al set.^9^ The remaining sites are likely novel.

Each entry was annotated with a pathogenicity tag, which we sorted into three bins: “pathogenic,” “benign,” and “unspecified.” The proportions of these tags in the set of all exonic SNPs, all C>U SNPs, and all predicted editing sites were calculated (Figure 2C and Figure 2D). Of the changes in the C>U editing set, 19,285 (19.0%) were tagged as pathogenic, 9459 (9.3%) were tagged as benign, and 72,821 (71.7%) were unspecified. The percentage of pathogenic SNPs were higher in the RNAsee predicted set; in this set, 1046 (22.7%) of SNPs were tagged as pathogenic, 3132 (68.1%) as unspecified, and 422 (9.2%) as benign. This also meant that a higher percentage of pathogenic C>U sites were predicted editing sites (5.4%) than unspecified (4.3%) or benign (4.5%). The percentage of benign SNPs was relatively consistent between the sets, but the percentage of pathogenic sites was highest in the union set and lowest in the set of all exonic SNPs.

### Associations with disease

Of 101,565 sites in the all C>U set, 72917 (71.8%) were associated with at least one disease, condition, or phenotype in ClinVar. To determine which areas of health are most likely to be affected by APOBEC3A/G-mediated RNA editing, we decided to survey these conditions.

We labelled each SNP with the MeSH subject headings that most closely matched its associated conditions. In total, there were 6,534 unique conditions associated with C>U SNPs. 575 conditions (8.8%) could not be matched with a good single MeSH equivalent and were labelled “Not found.” In total, 7181 SNPs (9.3%) were associated with “Not found” at least once, including 401 SNPs considered pathogenic.

The top-level subject headings corresponding to the initial MeSH labels were found for each SNP, with each top-level term labelling an SNP no more than once. We counted the number of times each top-level subject heading or “Not found” was associated with a SNP in the all C>U and RNAsee predicted sets. Because this work is focused on the potential effects of RNA editing on human health, we focused on those SNPs labelled pathogenic. The top ten most common top-level MeSH subject headings associated with pathogenic SNPs in the all C>U and RNAsee predicted sets are shown in Figures 3A and 3B, respectively.

**Figure 3.**
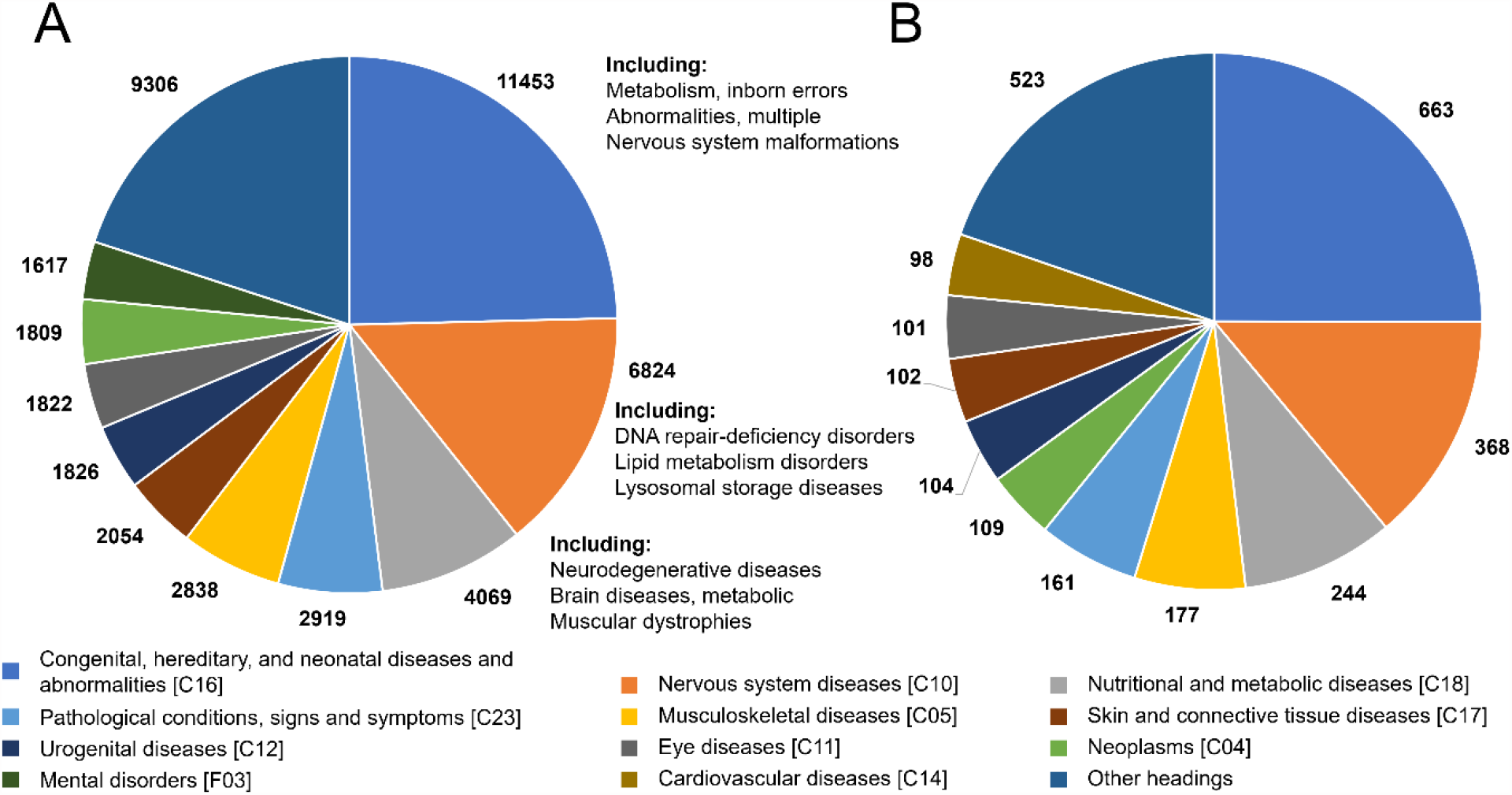
Potential pathogenic APOBEC3A/G targets are associated with similar diseases to the set of all C>U SNPs. The number of times the top ten most common top-level MeSH subject headings are associated with the set of all pathogenic C>U SNPs (A) and the pathogenic RNAsee predicted set (B) are shown. The most common types of conditions are similar between the two groups, with congenital abnormalities, nervous system diseases, and nutritional and metabolic diseases being the most common types of conditions associated with both sets.

In both sets, the most common associated heading was “Congenital, Hereditary, and Neonatal Diseases and Abnormalities.” Because ClinVar pathogenic variations are thought to be causally linked to their associated conditions, all conditions associated with these variations should either be congenital or a result of a de novo tissue mutation, for instance, in a neoplasm. “Nervous System Diseases” and “Nutritional and Metabolic Diseases” were the second and third most common subject headings in both groups, which may mean C>U RNA editing is most likely to negatively influence human health in these two areas. The proportions of most subject headings were similar between the two groups, with only slight differences in their relative rankings.

To further elucidate the potential effects of APOBEC3-mediated RNA editing on human health, we wanted to examine the proportion of diseases with any possible influence from APOBEC3-mediated RNA editing. To do this, we counted the number of third-level MeSH headings (grandchildren of top-level headings) with at least one predicted editing site. To find an upper bound on the number of conditions affected, we also counted the number associated with at least one C>U SNP (Figure 4).

**Figure 4.**
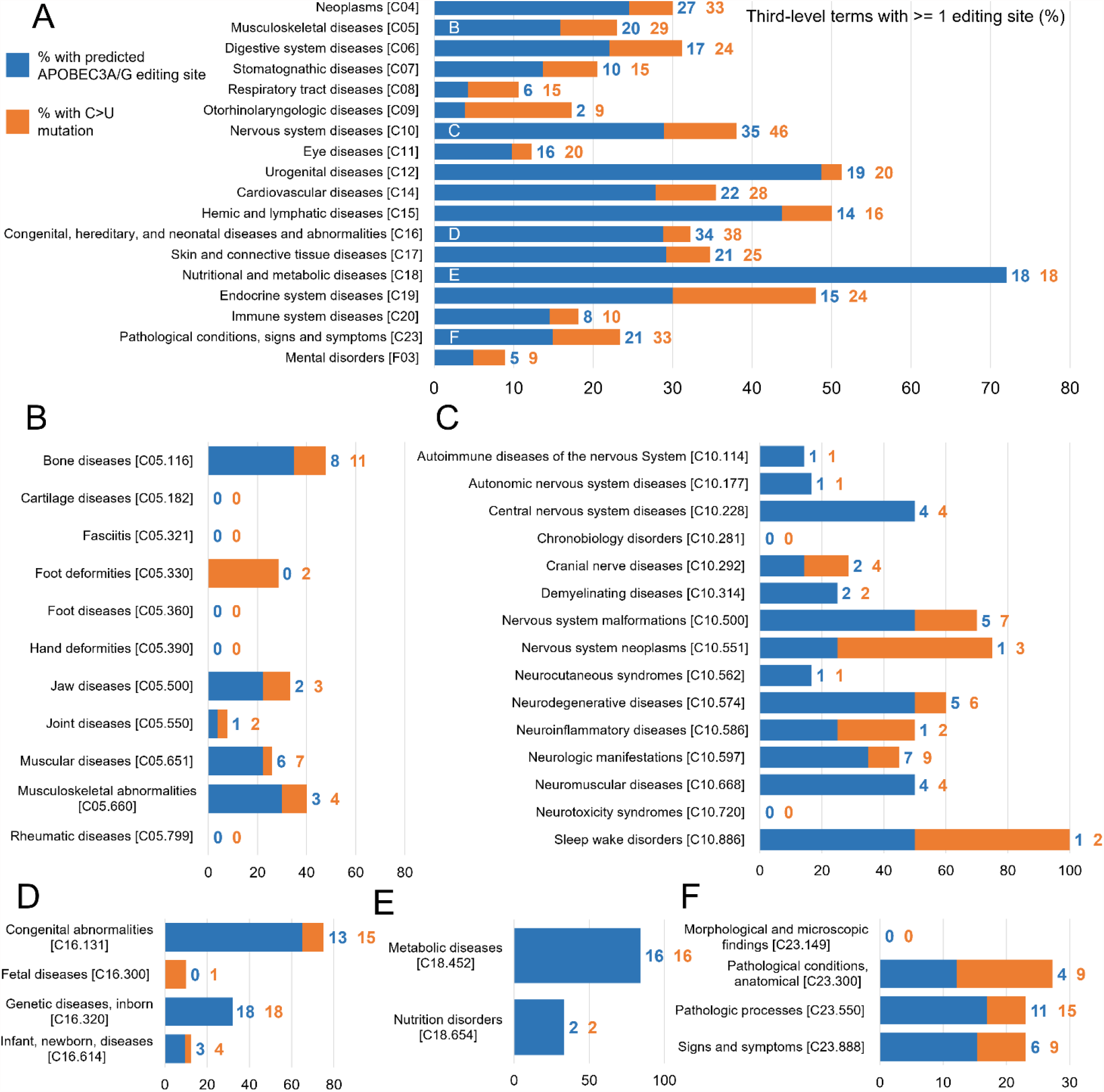
Proportion of conditions associated with at least one potential APOBEC-mediated RNA editing site. (A) For select top-level MeSH terms, the number of grandchild terms with at least one predicted APOBEC3A/G editing site was calculated (blue). In addition, the number of grandchild terms with at least one associated C>U mutation, but no predicted editing site, was found (orange). This was divided by the total number of grandchild terms for each top-level term to approximate the percentage of diseases associated with each top-level term that could be influenced by C>U RNA editing at a known site of genetic variation. The total number of terms counted can be found at the end of each bar, and the tree number of each term can be found in brackets at the end of its label. Additional graphs, showing the percent of child terms for each second-level term, are provided for the child terms of the top five most common top-level terms: (B) Musculoskeletal Diseases, (C) Nervous System Diseases, (D) Congenital, Hereditary, and Neonatal Diseases and Abnormalities, (E) Nutritional and Metabolic Diseases, and (F) Pathological Conditions, Signs and Symptoms. C>U RNA editing may play a role in at least one disease for almost every category of non-acquired human disease.

Almost all top-level types of non-traumatic, non-infectious conditions recognized by MeSH had at least one grandchild term associated with at least one predicted editing site. Some types of diseases had as few as 4% of their grandchild terms with an associated editing site (otolaryngologic diseases, respiratory tract diseases), whereas 72% of third-level nutritional and metabolic diseases were associated with predicted editing sites. Notably, some top-level headings with a large number predicted editing sites had a lower percentage of diseases with associated predicted editing sites.

## Discussion

We searched a list of known non-synonymous DNA SNPs to see which of those polymorphisms, and the resultant variant proteins, could also be caused by APOBEC3A/G-mediated RNA editing. We found that about 4.5% of known SNPs which result in C>U mRNA changes are also potential APOBEC3A/G editing sites. If APOBEC3A/G RNA editing regularly occurs at even a fraction of these sites, this could result in meaningful effects on human health. It has previously been demonstrated that interferon-rich environments, such as inflamed tissues, increase APOBEC3-mediated RNA editing in a variety of cell types.^8,12-15^ Transient increases in RNA editing could result in the production of variant proteins, which could affect recovery time and result in sequelae following periods of inflammation.

Over 40% of exonic, non-synonymous human SNPs recorded in the ClinVar database are associated with a C>T change on one strand. This is far more than expected by random chance alone; counting the number of non-synonymous single nucleotide changes a codon can undergo, only 14.4% are C>T or G>A. This high percentage may be due, in part, to rare uncorrected C>U DNA editing events catalyzed by enzymes such as APOBEC3A/G and AID (activation-induced cytidine deaminase).^8,10^ APOBEC3A was already known to extensively edit DNA in neoplasms, which lends credence to this theory.^10^ Interestingly, neoplasms were only the sixth most common subject heading associated with editing sites predicted by RNAsee. Therefore, APOBEC3A/G DNA editing is likely a factor in pathogenic SNPs and the creation of variant proteins beyond its known activity in neoplasms. RNA editing could have a likewise unknown role in health.

When the union method of RNAsee was benchmarked, it returned about 3.2% of cytosines in the proportional set as potential APOBEC3A/G editing sites, including false positives. However, when run on C>U sites in the ClinVar database, it identified 4.5% (4600) of C>U SNPs and 5.4% (1046) of pathogenic C>U SNPs as potential editing sites. This higher percentage supports the idea that some of the C>T and G>A polymorphisms recorded in ClinVar result from APOBEC3A/G DNA editing, leading to a sample that was biased towards higher APOBEC3A/G editing affinity. It was previously found that APOBEC3A was more active against RNA than DNA in cancer cells, and this activity occurred at similar sites in both.^23^ Therefore, if many of these SNPs were indeed originally caused by APOBEC3A/G ssDNA editing, RNA editing likely occurs at the same sites.

When compared to SNPs associated with C>T changes on one strand, sites associated with C>U changes in RNA were underrepresented. By random chance, half of C>T and G>A changes should be on the correct strand to result in C>U RNA changes, but only 16.5% of SNPs actually resulted in C>U changes. This could be explained if the enzymes which catalyze C>U deamination have easier access to or higher affinity for a DNA strand and segment that is being transcribed. This could also result from evolutionary pressure against the appearance of good C>U editing sites in RNA if C>U RNA editing events are generally deleterious.

We believe that these results, alongside existing evidence, support the significance of APOBEC3A/G RNA editing to human health (Figure 5). Strong evidence suggests that expression of APOBEC3 enzymes increase in certain tissues when exposed to increased environmental interferons, as in the case of tissue inflammation.^8,12-15^ These periods of increased enzyme expression provide increased opportunities for APOBEC3A/G-mediated mRNA editing events to occur, particularly in optimal stem-loop structures.^8,10^

**Figure 5.**
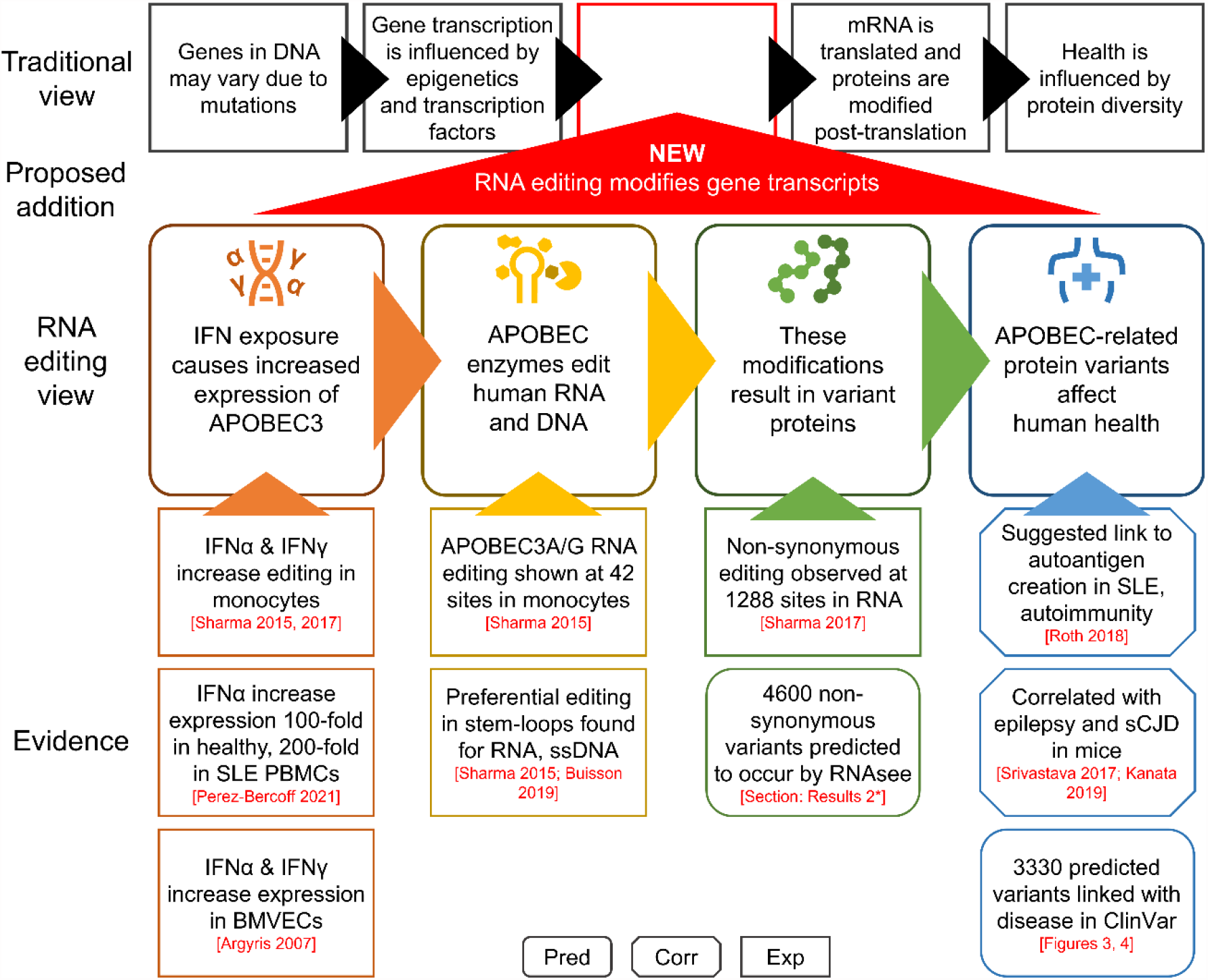
Selected evidence for the influence of RNA editing on human health. In the traditional model of biology, protein diversity results from mutations in DNA and post-translational modifications, and the expression of proteins is regulated via epigenetics and transcription factors. We propose adding RNA editing to this view. Current evidence suggests that APOBEC family enzymes are expressed when some cell types are exposed to environmental IFN. Some APOBEC enzymes, such as APOBEC3A and APOBEC3G, have been shown to edit human RNA. This editing has been shown to cause non-synonymous changes in mRNA, and we used RNAsee to predict additional sites that may also undergo editing. The effects of this editing on human health are still under researched, but some prior work has suggested links between APOBEC-mediated RNA editing and certain autoimmune and neurological disease. Our predictions additionally suggest that at least one potential editing site is associated with most disease categories recognized in MeSH subject headings. It is essential that future work attempts to investigate and confirm or refute the links between RNA editing and human health as proposed in this and other works. *The box shape of evidence boxes represents the type of study the information is taken from. Rounded edges are computational (pred)ictions, cut corners are (corr)elational studies, and square corners are (exp)erimental studies.* *\* Results 2: Identification of possible editing sites* *PBMC: peripheral blood mononuclear cell; BMVEC: brain microvascular endothelial cells; IFN: interferon; SLE: systemic lupus erythematosus; sCJD: sporadic Creutzfeldt-Jakob disease*

These editing events have previously been shown to occur at locations which result in variant proteins due to missense or nonsense mutations.^12^ Beyond these known sites, we have found that over 40% of non-synonymous SNPs in the ClinVar database are associated with a C>T change on one DNA strand, much higher than expected by random chance and possibly indicative that C>U ssDNA editing has contributed greatly to the set of known human SNPs (Figure 1). Since similar substrates are targeted by APOBEC3A/G enzymes in DNA and RNA, and similar sites are edited in RNA and DNA, this suggests more frequent RNA editing is likely occurring at some or all of the 16% of non-synonymous SNPs associated with C>U RNA changes.^2,10,23^ RNAsee specifically suggested 4600 of these sites, or 4.5% of all C>U SNPs, as particularly probable editing sites, and 62 of those sites are known editing sites included in the Asaoka et al dataset.^9^

The ultimate question is: does this editing actually affect human health? Our work suggests it does. Of those sites suggested as probable RNA editing sites by RNAsee, 1046 were tagged as pathogenic, and an additional 3132 have unknown or unspecified pathogenicity. Over 20% of third-level MeSH subject headings associated with nutritional and metabolic, neoplastic, cardiovascular, and nervous system diseases were associated with at least one predicted editing site. In addition, more studies are finding links between RNA editing and human disease. For instance, APOBEC-mediated editing has been correlated with epilepsy and sCJD in mouse models.^16,17^ One study on the linkage between SLE and RNA editing even suggested RNA editing as a mechanism for autoantigen formation in autoimmune diseases.^18^ Therefore, there is a high likelihood that RNA editing affects human health in some or all of the areas of disease noted in this study or existing studies on RNA editing, particularly when there is an environmental pressure causing increased APOBEC3 activity.

This paper only considers the activity of the APOBEC3A/G enzymes. If APOBEC3A and APOBEC3G could cause polymorphisms at some of the loci identified in this paper and mediate some of the disorders noted, it seems likely that other APOBEC or ADAR-family enzymes are simultaneously editing RNA at entirely distinct sites, activated by distinct conditions, and with distinct effects on health. Therefore, to fully explore this mechanism behind protein diversity, future research should be devoted to (1) describing the optimal substrates for RNA editing enzymes, (2) finding the conditions in which these enzymes are most active, and (3) identifying the effects of variant RNA or proteins created by RNA editing on protein diversity and human health. In this way, we can add to our basic understanding of human molecular biology.

## Conclusion

Classically, our biodiversity is thought to come from our constitutive genetics, epigenetic phenomenon, transcriptional differences, and post-translational modification of proteins. Here, we have shown evidence that RNA editing, often stimulated by environmental factors, could account for a significant degree of the protein biodiversity leading to human disease. In an era where worries about our changing environment are ever increasing, from the warming of our climate to the emergence of new diseases to the infiltration of microplastics and pollutants into our bodies, understanding how environmentally sensitive mechanisms like RNA editing affect our own cells is essential. Future research will apply our analysis to the transcriptomes and proteomes of specific disease conditions in order to further clarify the functional and predictive role of APOBEC-mediated C>U RNA editing in human disease.

## Methods

### Extraction of disease variant data

The ClinVar database, maintained by the National Center for Biotechnology Information (NCBI), collects information regarding DNA variants found in patient samples, including assertions of clinical significance and evidence to support those assertions.^1^ We extracted the annotated set of DNA mutations from ClinVar on May 18, 2022 via FTP.^2^ All variants which were not SNPs (“Type” of “single nucleotide variant”) were excluded. Two records were provided for each variant corresponding to the GRCh37 and GRCh38 reference genomes. Records corresponding to GRCh37 were excluded. Finally, because this study was intended to examine RNA editing as a potential source of protein diversity in human cells, all DNA variants which would not result in protein variants (including non-exonic or exonic but synonymous variants) were excluded.

For each SNP, we extracted the following information: the gene, allele ID, any amino acid change resulting from the mutation, assertion of clinical significance, rsID, original and mutant nucleotides, and list of phenotypes associated with the variant. To get a baseline for the frequency of different SNPs, we used the original and mutant nucleotides (found in the “ReferenceAlleleVCF” and “AlternateAlleleVCF” columns) for each SNP. Entries with non-specific nucleotides (e.g. A>X) or like-to-like nucleotide changes (e.g. A>A) were excluded.

Entries without any associated amino acid change recorded were also excluded. We also binned the clinical significance into three categories (“pathogenic,” “benign,” or “unspecified”) based on whether the significance tag contained the words “pathogenic,” “benign,” or neither.

Coding sequence files were obtained from the CCDS.^3^ An attempt was made to download a human coding sequence file for any gene with at least one SNP which matched the inclusion criteria. If said file did not exist or was incompatible with the data contained within ClinVar (i.e. the original amino acid in the record is not coded for by the codon at the corresponding location in the coding sequence file), variants in that gene were excluded. Out of 622,222 SNPs matching the inclusion criteria, 617,363 exonic single nucleotide polymorphisms across 9228 genes were analyzed. The amino acid change recorded in each entry and the corresponding codon in the coding sequence file were used to determine whether each SNP could be associated with C>U RNA editing. Those DNA polymorphisms that resulted in a C>U change in the transcribed mRNA were included in the C>U set and analyzed using RNAsee. In total, 101,565 sites were assessed for their likelihood of being APOBEC3A/G editing sites.

### RNAsee

RNAsee is a publicly available Python package. It can be found at https://github.com/ram-compbio/RNAsee. We previously developed RNAsee to identify potential APOBEC3A/G editing sites. RNAsee v1 was a rules-based method that used a simple stem-identification algorithm and a minimum free energy score assigned by ViennaRNA to rank cytosines within a gene from most to least likely to undergo RNA editing.^4,5^ During the development of RNAsee v2, we decided to utilize the set of known APOBEC3-mediated RNA editing sites published in Asaoka et al. to both improve and benchmark RNAsee.^6^ All cytosines identified in Asaoka et al. were considered editing sites, and all other cytosines in the same genes were considered non-editing sites.^6^

We refined the rules-based model to score sites based on sequential features commonly observed in this dataset in addition to the presence and strength of a stem-loop structure. We developed a novel scoring algorithm, and we trained a scoring threshold for the rules-based model for best F1 score on a subset of the dataset.

We also added a machine learning model to RNAsee. Four types of classification models were initially considered: support vector machines, logistic regression, decision tree, and random forest. We also considered different methods of vectorizing the sequence surrounding the potential editing site based on number of nucleotides included (a 25- or 15-nucleotide stretch surrounding the site) and method of encoding nucleotides (4 or 2 binary integers per nucleotide). Finally, we attempted to accommodate the highly imbalanced class sizes between editing and non-editing sites (1:468) by downsampling non-editing sites and/or using SMOTE-based up-sampling of editing sites. Ultimately, based on initial benchmarking performance and area under the receiver operator curve (AUROC) figures, we chose a random forest model that takes as input a 25-nucleotide stretch surrounding a cytosine, vectorized into 2 binary integers per nucleotide (isPurine and pairsGC), trained on a set of sites downsampled for non-editing sites and with an increased proportion of non-editing sites that received high scores when scored by the rules-based model

RNAsee v2 includes two primary methods and two consensus methods for editing site identification:

- **Rules-based model**. The rules-based model assigns a score to cytosines if, and only if, they are found at the 5’ end of a 3-4 nucleotide loop in a stem-loop structure. Scores are calculated based on the strength of the stem (3*(the number of GC pairs) + (the number of AU pairs)) and the presence or absence of specific sequential features (+2 for a uracil in the loop or a purine following the cytosine and −2 for a guanine in the loop). A site is considered an editing site if it receives a score of greater than nine.
- **Random forest model**. The random forest model was created as an instance of the RandomForestClassifier class from scikit-learn.^7^ The model takes as input a 50-bit vector representing 15 nucleotides preceding and 10 following a cytosine. Every nucleotide is represented by 2 bits; one for whether that nucleotide is a purine and another for whether it can participate in GC pairing. The model was trained and tested using a 70-30 training/testing split of known editing and non-editing istes. A site is considered an editing site if its probability of being an editing site is over 0.5.
- **Consensus models**. The intersection and union models combine the outputs of the two primary methods using simple set operations. The intersection method outputs only sites returned by both primary models, and the union method outputs all sites returned by either primary model.

### Performance benchmark of RNAsee

Two datasets were used to benchmark RNAsee: the testing set and the proportional set. The former included all sites in the random forest model’s testing set. It contained a 3:1 ratio of non-editing to editing sites. Because this ratio is greatly different from that found in the whole Asaoka et al. dataset, we also created the proportional set. This set contains all sites found in the testing set, plus additional non-editing sites to bring the ratio of non-editing sites to editing sites to 468:1, similar to the original ratio.

During benchmarking, each model was used to predict which cytosines in both datasets are APOBEC3 editing sites. The number of non-editing and editing sites in each model’s predictions were tallied and used to calculate metrics including sensitivity and precision. For the primary models, AUROC figures were also calculated.

Because this paper focused on fully exploring the potential effects of APOBEC3A/G-mediated RNA editing on human health, we decided to use the union model, which showed the highest sensitivity (82.8%). For each SNP in the C>U set, the original cytosine in question and its sequential context was passed as input to the union model. All sites returned by the union model were considered potential APOBEC3A/G editing sites.

### Association with MeSH subject headings

The diseases and phenotypes associated with each site in the C>U set were extracted. All unique conditions were extracted into a single list. For each term in this list, the most relevant MeSH subject heading (descriptor or supplemental concept record) was found. MeSH mappings were used to associated supplemental concept records with tree terms unless a mapping contained a qualifier that suggested the supplemental concept record was not a child of the mapped term (such as “abnormalities”). Each term was then associated with the chosen subject heading and every parent subject heading up to the top level. If no single subject heading encapsulated the given term, that term was associated with the phrase “Not found.” “Not found” was treated as a top-level subject heading for analysis purposes. Finally, the terms were re-associated with the SNPs.

The number of times each subject heading appeared across all SNPs was counted. Each subject heading was counted up to once per SNP, even if that SNP was associated with multiple conditions corresponding to that subject heading. Using the same method and same list of condition-subject heading associations, the number of times each subject heading appeared across the union set was also calculated. Additional counts were also generated for the subsets of pathogenic, benign, or unspecified SNPs in each set. Analysis was primarily carried out on the counts of top-level subject headings.

Additional figures were calculated for the percentage of diseases with at least one C>U SNP and/or editing site associated. For the purposes of this work, the third level of MeSH subject headings (grandchildren of top-level terms) were considered to represent individual diseases. For each top-level term and select second-level terms, the number of associated third-level terms was counted. If at least one C>U SNP or editing site mapped directly to a third-level term or to child terms of that third-level term, the term was counted as having a C>U SNP or editing site. The percentages of diseases with at least one associated C>U SNP and diseases with at least one editing site were then calculated by dividing these counts by the total number of third-level terms.

## Declarations

The authors have no conflicts of interest to report.

## Acknowledgements

This work has been supported in part by grants from NIH NLM [T15LM012495, 1R25LM014213], NIAAA [R21AA026954, R33AA0226954], NIDA [1K01DA056690], NIST [60NANB22D168], and NCATS [UL1TR001412]. This study was funded in part by the Department of Veterans Affairs. The authors would like to acknowledge the Center for Computational Research (CCR) at University at Buffalo for computational support. We would also like to thank all members of the Samudrala Computational Biology Group.

## Notes

https://github.com/ram-compbio/RNAsee

### Competing Interest Statement

The authors have declared no competing interest.

